# Improving Gene Regulatory Network Inference by Incorporating Rates of Transcriptional Changes

**DOI:** 10.1101/093807

**Authors:** Jigar S. Desai, Ryan C. Sartor, Lovely Mae Lawas, SV Krishna Jagadish, Colleen J. Doherty

## Abstract

Organisms respond to changes in their environment through transcriptional regulatory networks (TRNs). The regulatory hierarchy of these networks can be inferred from expression data. Computational approaches to identify TRNs can be applied in any species where quality RNA can be acquired, However, ChIP-Seq and similar validation methods are challenging to employ in non-model species. Improving the accuracy of computational inference methods can significantly reduce the cost and time of subsequent validation experiments. We have developed ExRANGES, an approach that improves the ability to computationally infer TRN from time series expression data. ExRANGES utilizes both the rate of change in expression and the absolute expression level to identify TRN connections. We evaluated ExRANGES in five data sets from different model systems. ExRANGES improved the identification of experimentally validated transcription factor targets for all species tested, even in unevenly spaced and sparse data sets. This improved ability to predict known regulator-target relationships enhances the utility of network inference approaches in non-model species where experimental validation is challenging. We integrated ExRANGES with two different network construction approaches and it has been implemented as an R package available here: http://github.com/DohertyLab/ExRANGES. **To install the package type:** devtools::install_github(“DohertyLab/ExRANGES”)

## Introduction

Transcriptional regulatory networks (TRN) provide a framework for understanding how signals propagate through a molecular network and result in transcriptomic changes. These regulatory networks are biological computational modules that carry out decision-making processes and, in many cases, determine the ultimate response of an organism to a stimulus ^1^. Understanding these regulatory networks provides access points to modulate these responses through breeding or genetic modifications. The first step in constructing such networks is to identify the primary relationships between regulators such as transcription factors (TFs) and the target genes they control.

Experimental approaches such as Chromatin Immunoprecipitation followed by sequencing (ChIP-Seq) can identify direct targets of transcriptional regulators. However, ChIP-Seq must be optimized to each specific TF and antibodies must be developed that recognize either the native TF or a tagged version of the protein. This can present a technical challenge particularly for TFs where the tag interferes with function, for species that are not easily transformable, or for tissues that are limited in availability ^2^. Since global transcript levels are comparatively easy to measure in most species and tissues, several approaches have been developed to identify connections between regulators and their targets by examining the changes in transcription levels across many samples ^3–6^. These inferred approaches can provide a first approximation of regulatory interactions that can be used to guide experimental approaches. The assumption of these approaches is that the regulatory relationship between a regulator TF and its targets can be discerned from a correspondence between the RNA levels of the regulator gene and its targets. If this is true, then given sufficient variation in expression, the targets of a given factor can be predicted based on associated changes in expression. Initial approaches designed to do this focused on the correlation between regulators and targets, assuming that activators are positively correlated and repressors are negatively correlated with their target expression levels ^7^. For almost two decades, these approaches successfully identified relationships between regulators and targets. Updates to this simple idea have included pre-clustering of transcript data, modifying regression analysis, incorporating training classifier models, and incorporating prior biological knowledge or additional experimental data. Each of these has improved the ability to identify connections between regulators and targets, even in sparse and noisy data sets ^4–6, 8–11^. For microorganisms, substantial experimental data identifying TF binding locations and the transcriptional response to TF deletions is available and has been organized into efficient databases ^12–14^. This approach has enabled the prediction of TRN from expression data not only in unique conditions in the model species where the data was generated, but has also been extended to predict TF-target gene relationships in homologous species ^15–19^. In 2010, the DREAM5 challenge evaluated the ability of different methods to identify TRN from gene expression data sets ^10^. One of the top performing methods was GENIE3 ^8^. This method uses the machine learning capabilities of random forest to identify targets for selected regulators ^20, 21^. Other successfully implemented approaches include SVM ^22^, CLR ^6^, CSI ^23, 24^, ARACNE ^5^, Inferelator ^4^, and DELDBN ^9^. Common to these methods is the use of transcript abundance levels to evaluate the relationship between a regulator and its putative targets. Experiments performed in time series can provide additional kinetic information useful for associating regulators and targets. Many approaches have been developed that take advantage of the additional information available from time series data as reviewed in ^25, 26^. However, the steady-state transcript level as measured by most high-throughput transcriptional assays such as RNA-Seq is a measure of both transcriptional activity and mRNA stability. Therefore, the correlation between expression levels alone may not provide a direct assessment of transcriptional regulation as it can be confounded by the RNA stability of the target. Further complicating the identification of regulator relationships is the fact that a single gene can be regulated by different transcription factors in response to different stimuli.

Here we present an approach that extends current approaches to TRN construction by emphasizing the relationship between regulator and targets at the time points where there is a significant change in the rate of expression. We demonstrate that: 1) Focusing on the rate of change captured previously unrecognized characteristics in the data, identifying experimentally validated regulatory relationships not detected by the standard approaches. 2) Combining expression level and the rate of change results in an improved identification of experimentally validated regulatory relationships.

We first evaluate the significance of the rate changes at each consecutive time point on a per-gene basis: RANGES (RAte Normalized in a GEne Specific manner). We then combined the expression level and significance of this rate change in ExRANGES (Expression by RANGES) to prioritize the correlation between regulators and targets at time points where there is a significant change in gene expression. ExRANGES improved the ability to identify experimentally validated TF targets in microarray and RNA-Seq data sets across multiple experimental designs, and in several different species. We demonstrate that this approach improves the identification of experimentally validated TF targets using GENIE3 ^8^, and anticipate that it will offer a similar benefit to when combined with other network inference algorithms.

## Results

### ExRANGES Improves Identification of Circadian TF Targets in a Circadian Data Set

The assumption behind using correlation in gene expression to identify relationships between TFs and their targets is that there is a predictable relationship between the expression of the TF regulator and its corresponding targets. For transcriptional activators, the target will accumulate as the TF regulator accumulates. Conversely, targets of repressors will decrease in expression as the repressor TF increases. Current approaches evaluate the correspondence in expression between the regulator TF and targets across all time points equally (hereinafter referred to as EXPRESSION). We developed ExRANGES, a method that adjusts the expression level based on how much that gene changes in expression in the following time step. Briefly, for each gene, we calculate the significance of each time step. The expression level is adjusted by this significance factor so that the expression level preceding a major change in expression is emphasized (Supplemental Fig. 1). We tested whether incorporating the rate of change via ExRANGES improves the overall ability to identify experimentally validated regulatory relationships. To evaluate the ability of the ExRANGES or standard EXPRESSION approaches to correctly identify targets of the TFs, we applied both approaches to the CircaDB data ^27^ (for description, see Supplemental Materials and Methods) using GENIE3. We compared the results of each approach to the targets identified experimentally using ChIP-Seq for five TFs involved in circadian regulation: PER1, CLOCK, NPAS2, NR1d2, and ARNTL ^28, 29^. Targets identified by each computational approach that were also considered significant targets in these published ChIP-Seq experiments were scored as true positive results. We calculated the ROC AUC for the five circadian TFs to compare the identification of true targets attained with GENIE3 using EXPRESSION values to the combination of expression and p-values using ExRANGES. We observed that for all five TFs, ExRANGES improved the identification of ChIP-Seq validated targets (Fig. 1A). Incorporation of a delay between regulator expression and target expression has previously been shown to improve the ability to identify regulatory networks ^30^. A modification of GENIE3 incorporates this approach to identify transcriptional changes in the regulator that precedes the effects on the target by a defined time step. We compared ExRANGES to this modified implementation of GENIE3 that includes the time delay step (Supplemental Fig. 2A). As previously reported, we observe that the time step delay improved target identification for some TFs, compared to EXPRESSION alone, although in this data set, target identification for CLOCK, PERI, and NR1D2 TFs did not improve. However, for all five TFs, ExRANGES outperformed both the EXPRESSION and time-delay approaches in identifying the true positive targets of each TF; although for CLOCK, this improvement was minimal.

**Figure 1:**
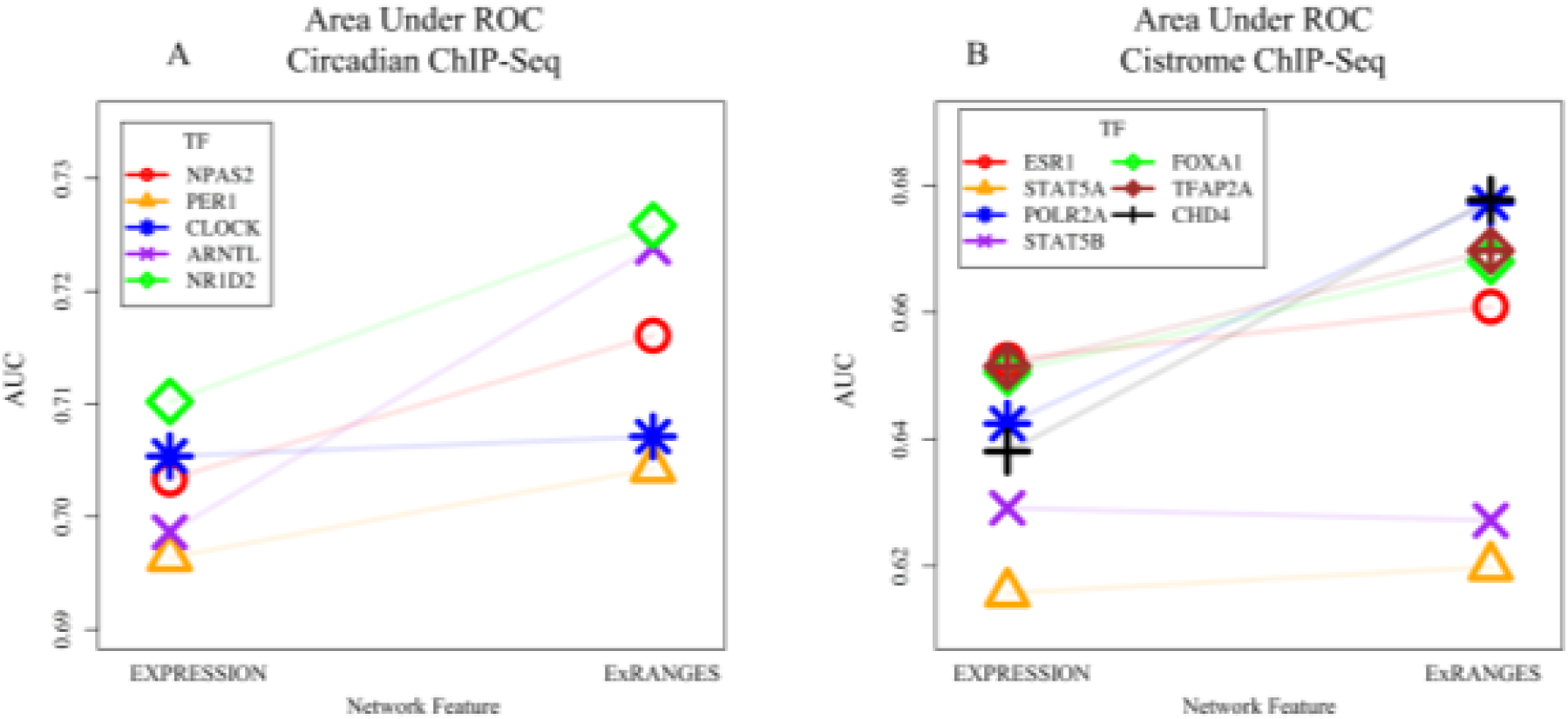
ExRANGES Outperforms EXPRESSION in Identifying Targets for Select TFs. A) ROC AUC for targets identified with GENIE3 using EXPRESSION or ExRANGES on five circadian TFs. The targets identified computationally were validated against ChIP-Seq identified targets ^28, 29^. B) ROC AUC for targets computationally identified by GENIE3 analysis using EXPRESSION or ExRANGES for seven TFs not known to be components of the circadian clock. Experimentally validated targets for these TFs were identified by ChIP-Seq in epithelial cells, a tissue not included in the expression data set ^31^.

### ExRANGES Improves Target Identification for TFs That Are Not Components of the Circadian Clock

To evaluate the performance of ExRANGES on TFs that are not core components of the circadian clock, we compared the ability to identify targets of additional TFs validated by ChIP-Seq. We selected seven TFs in our regulator list with ChIP-Seq data available from at least two experimental replicates performed in epithelial cells, a tissue not included in the CircaDB data set. The seven TFs are: ESR1, STAT5A, STAT5B, POL2A, FOXA1, TFAP2A, and CHD4 ^31^. Combining expression and rate change information using ExRANGES improved the AUC curve for five of the seven TFs (Fig. 1B, Supplemental Fig. 2B). As we observed above for CLOCK, STAT5A and STAT5B performed equally well but did not show significant improvement. STAT5A and STAT5B are known to be activated post-transcriptionally perhaps indicating why evaluating the change in expression of these TFs did not lead to improved target identification ^32–36^.

### ExRANGES Identified Targets have Less Variation Across the Time Series

The targets identified by ExRANGES or EXPRESSION approaches show moderate overlap in the ranked score of predicted targets (r^2^= 0.53); however, each network identifies different targets (Fig. 1 and Supplemental Fig. 3). To understand the difference in targets identified by EXPRESSION and ExRANGES we examined the variance in the expression levels for the top 1000 predicted targets of the 12 TFs identified by EXPRESSION or by ExRANGES across all 288 samples in the CircaDB data set. The targets identified by ExRANGES showed an overall lower coefficient of variation (CV) across all samples compared to targets identified by EXPRESSION (Fig. 2). The experimentally identified targets from ChIP-Seq showed low CV. The ability of ExRANGES to identify targets with lower CV than EXPRESSION may account for some of the improved identification of such the True Positive Targets.

**Figure 2:**
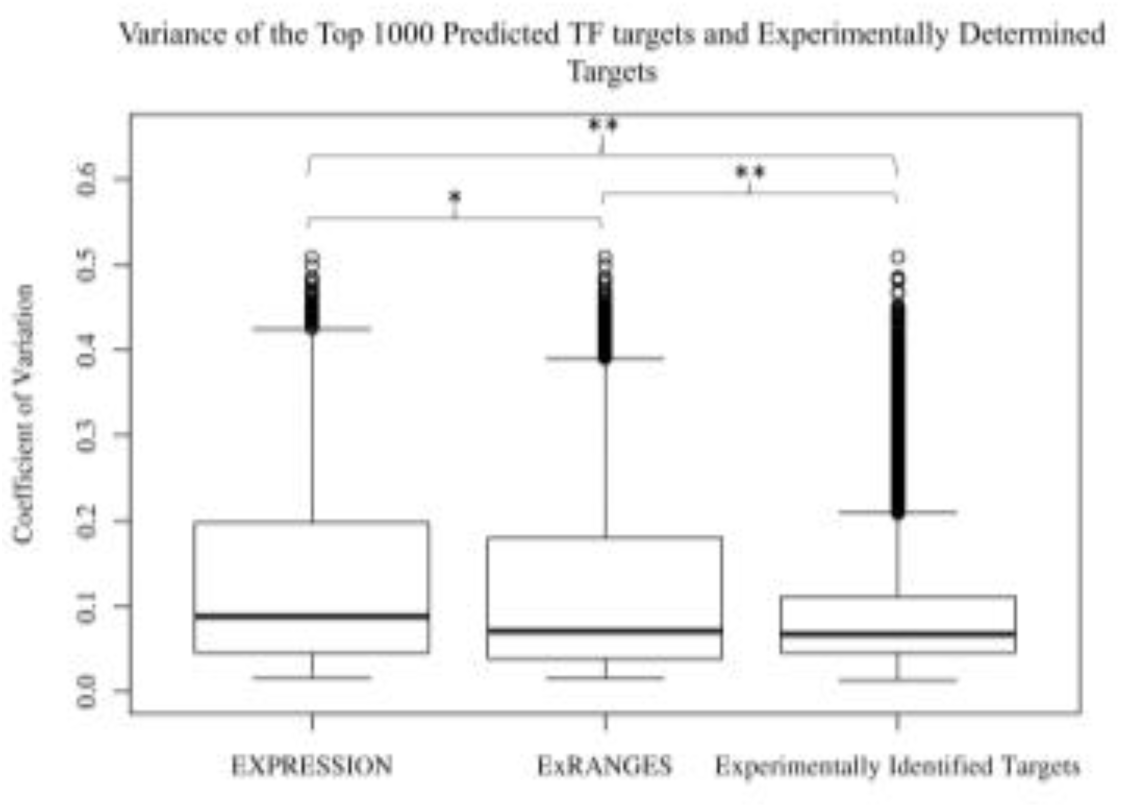
Targets Identified by ExRANGES and EXPRESSION have Different Variation across the CircaDB Data Set. Box plot showing the coefficient of variation (CV) for the expression levels of the top 1000 targets of each TF (ARNTL, CLOCK, NPAS2, NR1D2, PER1, ESR1, POL2A, FOXA1, TFAP2A, CHD4) predicted by GENIE3 using EXPRESSION or ExRANGES and the experimentally identified targets from ChIP-Seq. Targets identified using EXPRESSION show a greater expression CV across all samples compared to targets predicted with ExRANGES values. Experimentally determined targets showed the lowest CV (* *p*-value < 2.5e^-8^, ** *p*-value < 2e^-16^).

ExRANGES combines rate change and expression. To evaluate the contribution of the rate change component in the target identification, we generated a rate-based network using only the *p*-values of the rate change at each time step as our network feature. Using only rate change did not improve the overall identification of true positive targets (Supplemental Fig. 4). However, the targets identified in the rate-based network had lower overall variation in expression compared to the EXPRESSION identified targets. The CircaDB data consists of individual time series experiments from different tissues. Using rate change alone may enhance the identification of targets that have within tissue variation driven by changes across time compared to the larger overall variation between tissues observed in this data set. In contrast, EXPRESSION identified targets may favor those with large changes in expression between tissues. To evaluate how EXPRESSION and rate identified targets compared in variation within each time series in a single tissue versus between tissues, we compared the between tissue and within tissue standard deviation for the top 1000 targets identified by using EXPRESSION or rate change. The targets identified by EXPRESSION showed more variation between tissue types (Supplemental Fig. 5A). In contrast, the targets identified by rate change alone showed increased variation within each tissue time series compared to the EXPRESSION identified targets (Supplemental Fig. 5B).

We also compared the mean intensity level of the top 1000 predicted targets of the rate change and EXPRESSION approaches. We observed that the top 1000 targets of PER1 identified by EXPRESSION had higher intensity levels compared to the distribution of expression of all transcripts on the microarray (Supplemental Fig. 6A). In contrast, the top 1000 predicted targets of PER1 identified by rate change resembled the background distribution of intensity for all the transcripts on the array (Supplemental Fig. S6B). Likewise, the hybridization intensity of the genes identified as the top 1000 targets identified by EXPRESSION of all 1690 TFs considered as regulators was shifted higher compared to the background distribution levels (Supplemental Fig. 6C). The top 1000 targets of all 1690 TFs identified by rate change reflected the background distribution of hybridization intensity (Supplemental Fig. 6D). While hybridization intensity cannot directly be translated into expression levels, these observations suggest that there are features of the targets identified by rate change that are distinct from those identified by EXPRESSION.

### ExRANGES Improves Identification of TF Targets in Unevenly Spaced Time Series Data

Circadian and diel time series experiments are a rich resource providing temporal variance, which can be used to identify regulatory relationships. However, most available experimental data is not collected with this design. Often sample collection cannot be controlled precisely to attain evenly spaced time points. To evaluate the ability of ExRANGES to identify true targets of TFs across unevenly spaced and heterogeneous genotypes, we analyzed expression studies of viral infections in various individuals (“Respiratory Viral DREAM Challenge - Synapse ID syn5647810”; Liu *et al*. 2016) using both ExRANGES and EXPRESSION approaches. This data set consists of seven studies of blood samples from human patients. Multiple samples from an individual were taken over a seven to nine day period, depending on the specific study. Sampling was not evenly spaced between time points. In total 2372 samples were used, providing a background of 2231 consecutive time steps. Overall, the variance between samples was lower for this study than the circadian study examined above (Supplemental Fig. 7). The significance of a change in expression for each gene at each time step was compared to a background distribution of change in expression across all patients and time steps (2231 total slope changes). The targets identified using either EXPRESSION or ExRANGES were compared to ChIP-Seq identified targets of 83 TFs with available ChIP-Seq data from blood tissue ^31, 39^. We observed an overall improvement in the detection of ChIP-Seq identified targets for the 83 TFs with ExRANGES (Fig. 3A and B).

**Figure 3:**
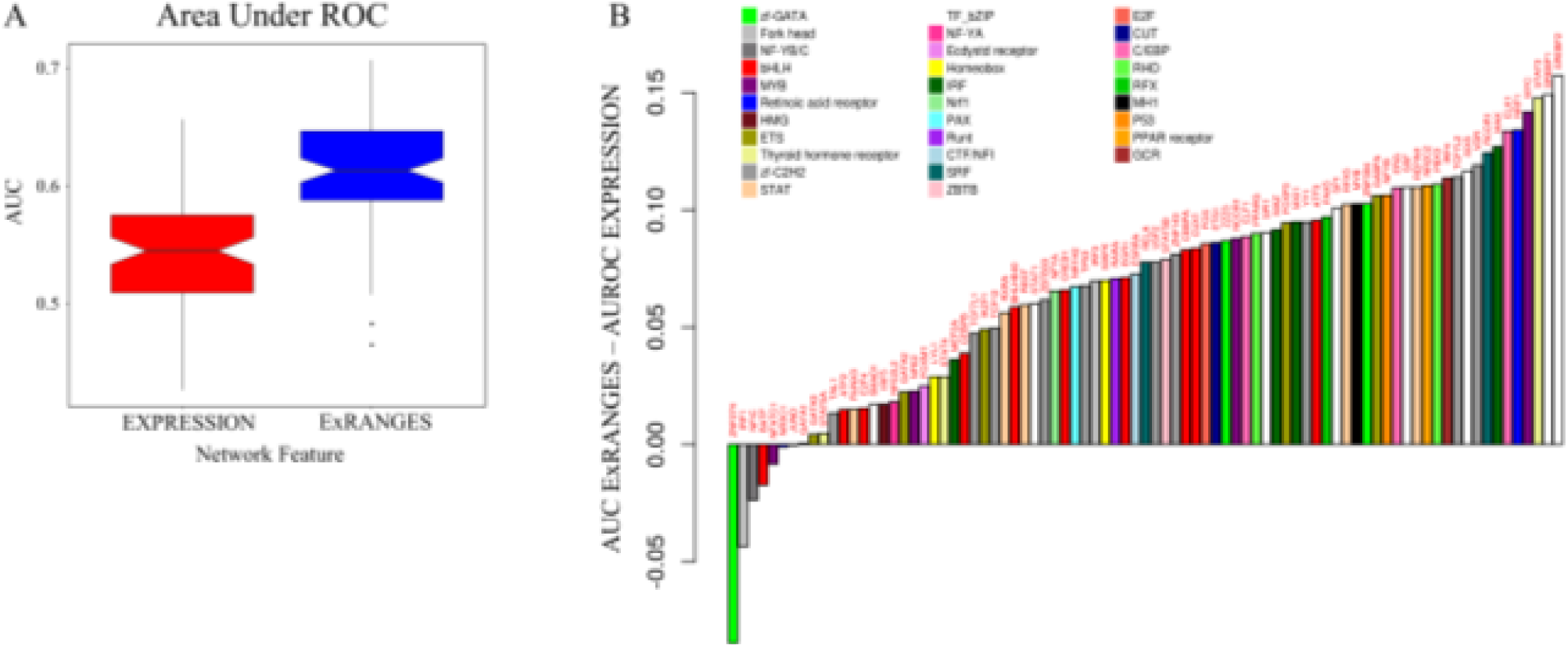
ExRANGES Improves Identification of Targets for most TFs from Unevenly Spaced Time Series Data. A) Box plot of ROC AUC for the GENIE3 analysis for all 83 TFs using either EXPRESSION or ExRANGES compared to ChIP-Seq identified targets. B) The difference between the ROC AUC of ExRANGES and EXPRESSION predicted targets is plotted individually for each of the 83 TFs tested, in ascending order. TFs are colored by TF family.

### ExRANGES Improves Functional Cohesion of Identified Targets

The true targets of a TF are likely to be involved in the same functional pathways, therefore functional enrichment can also be used to validate computationally identified TF targets ^40^. We compared the functional enrichment of the top 1000 targets predicted by either EXPRESSION or ExRANGES of the 930 TFs on the HGU133 microarray ^51^. The targets identified by ExRANGES for the majority of the TFs (590) showed improved functional enrichment compared to the targets identified by EXPRESSION (Fig. 4A and B). Likewise, when focusing on the 83 TFs with available ChIP-Seq data from blood, the majority of TF targets predicted by ExRANGES were more functionally cohesive compared to EXPRESSION targets as evaluated by GO slim (Fig. 4C). We observed that the improvement in the ranking of ExRANGES over EXPRESSION varies between the two validation approaches. For example, targets of the TF JUND identified by ExRANGES show no improvement over EXPRESSION when validated by ChIP-Seq identified targets, yet showed improved functional cohesion (Table ST1).

**Figure 4:**
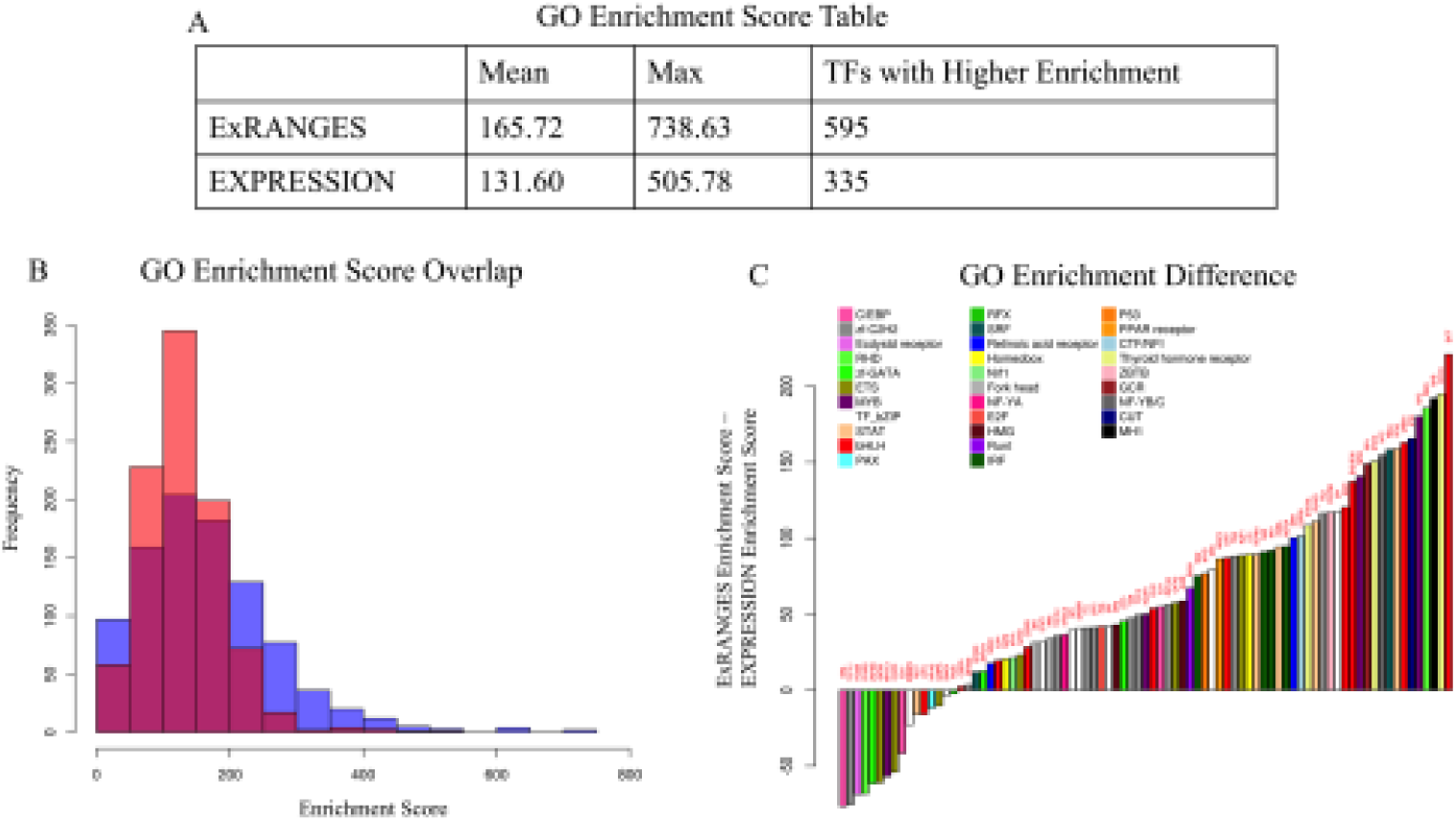
ExRANGES improves Functional Cohesion of Identified Targets. Gene Ontology term enrichment was calculated for the top 1000 predicted targets of 930 TFs using GENIE3 with either ExRANGES or EXPRESSION. Enrichment score is the sum of the –log10 of the *p*-value of each GO category. A) Summary of the enrichment scores for the top 1000 targets of all TFs on the microarray. B) The distribution of enrichments scores from EXPRESSION identified targets (red) and ExRANGES identified targets (blue). C) The difference in the enrichment score for the 83 TFs with available ChIP-Seq data (Fig 3). Positive values indicate TF targets with a higher enrichment score in ExRANGES compared to EXPRESSION.

### ExRANGES Improves TF Target Identification from RNA-Seq Data and Validated by Experimental Methods Other Than ChIP-Seq

To evaluate the performance of ExRANGES compared to EXPRESSION for RNA-Seq data we applied each approach to an RNA-Seq data set from *Saccharomyces cerevisiae* ^41^. This data set consisted of samples from six genotypes collected every fifteen minutes for six hours after transfer to media lacking phosphate. The slope background was calculated from 144 time steps. To evaluate the performance of ExRANGES compared to EXPRESSION approaches we calculated the AUC for the identified targets using GENIE3 for each of the 52 TFs using the TF targets identified by protein binding microarray analysis as the gold standard ^42^. For most TFs, the AUC was improved using ExRANGES (Fig. 5A).

**Figure 5:**
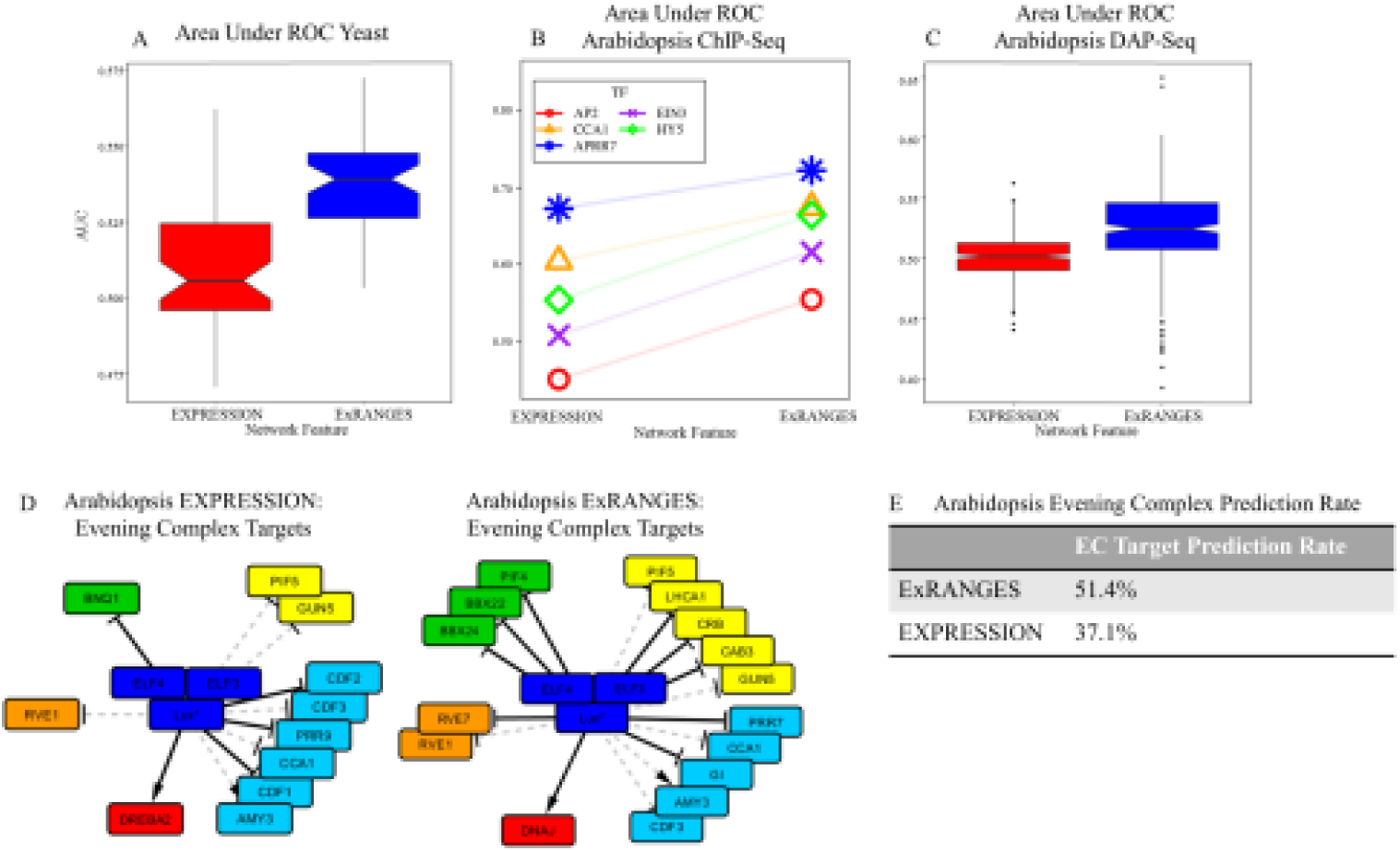
ExRANGES improves identification of TF targets validated by different methods. A) Box plots of the ROC AUC for targets identified for 52 yeast TFs by EXPRESSION or ExRANGES validated against experimentally identified targets from protein binding microarray data ^42^. B) ROC AUC for targets identified using GENIE3 with either EXPRESSION or ExRANGES for five Arabidopsis TFs validated against ChIP-Seq data. C). Box plot of AUC for targets identified for 307 Arabidopsis TFs by EXPRESSION and ExRANGES validated against DAP-Seq identified targets ^53^ D) Network of Arabidopsis Evening Complex Component (dark blue) regulated targets predicted with GENIE3 using either EXPRESSION or ExRANGES. Dashed edges are predicted targets that exist in both EXPRESSION and ExRANGES networks; solid edges are unique to the either EXPRESSION (left network) or ExRANGES (right network). The evening complex targets are colored by function: growth (green), photosynthesis (yellow), circadian (light blue), temperature-responsive genes (red), and light signaling (orange).* indicates the probeset corresponding to this gene can bind transcripts from more than one unique locus. E) Table of the prediction rate of evening complex targets of ExRANGES and EXPRESSION compared to those identified by ChIP-Seq ^56^.

We next evaluated the performance of EXPRESSION and ExRANGES on a set of data from Arabidopsis consisting of 144 samples collected every four hours for two days in 12 different growth conditions ^43–47^. Even though fewer ChIP-Seq data sets are available to validate the predicted targets in Arabidopsis, we were able to evaluate the performance of the algorithms for five TFs with available ChIP-Seq or ChIP-Chip identified targets performed in at least two replicates ^48–52^. We observed that for all five TFs, ExRANGES showed improved identification of the ChIP-based true positive TF targets (Fig. 5B). To evaluate a larger range of targets we compared our predicted targets by EXPRESSION or ExRANGES to 307 TFs targets identified by DAP-Seq ^53^. We observed that ExRANGES also showed an improved ability to identify targets as validated by DAP-Seq compared to EXPRESSION (Fig. 5C).

To evaluate the performance of ExRANGES compared to EXPRESSION, we constructed two networks of the Arabidopsis circadian clock; a TF-TF network of only the core clock components (Supplemental Fig. 8) and a TRN of the output from the Evening Complex (Fig. 5D and E) using GENIE3 with either ExRANGES or EXPRESSION as input. We limited our network of the core circadian clock to only genes previously established as associated with the circadian clock. The cyclic nature of circadian regulation makes modeling these interactions a challenge. ExRANGES correctly identifies more of the complex interactions within the morning loop that are supported by experimental data as reviewed in Greenham and McClung ^54^ compared to EXPRESSION (Supplemental Fig. 8). For example, ExRANGES correctly identifies interactions between CCA1/LHY and PRR9,7,5 that are not detected using EXPRESSION alone. The Evening Complex of the circadian clock controls the transcriptional regulation of many genes related to growth and light signaling responses ^55^. To evaluate the output of the circadian clock, we compared the targets of the Evening Complex proteins, ELF3, ELF4, and LUX, as identified by ChIP-Seq ^56^. The TRN constructed using ExRANGES identifies more of the evening complex targets than the TR N constructed using EXPRESSION as the input. In the top 10% of predicted interactions, more than 50% of the EC targets were called by ExRANGES and less than 40% was called by EXPRESSION (Fig. 5E). Indicating that the output of the Evening Complex is more reliably predicted using ExRANGES as an input to GENIE3.

### Application of ExRANGES to Smaller Data Sets with Limited Validation Resources

Time series data offers several advantages; however, it also increases the experimental costs. We have shown that using ExRANGES improves the performance of GENIE3 on large data sets as validated by ChIP-Seq (228 samples in mouse, 2372 in human, and 144 in arabidopsis) (Fig. 6). Since our interest is to develop a tool that can assist with the identification of regulatory networks in non-model species, we wanted to determine if ExRANGES could also improve identification of TF targets in sparsely sampled data sets where there is limited validation data available.

**Figure 6:**
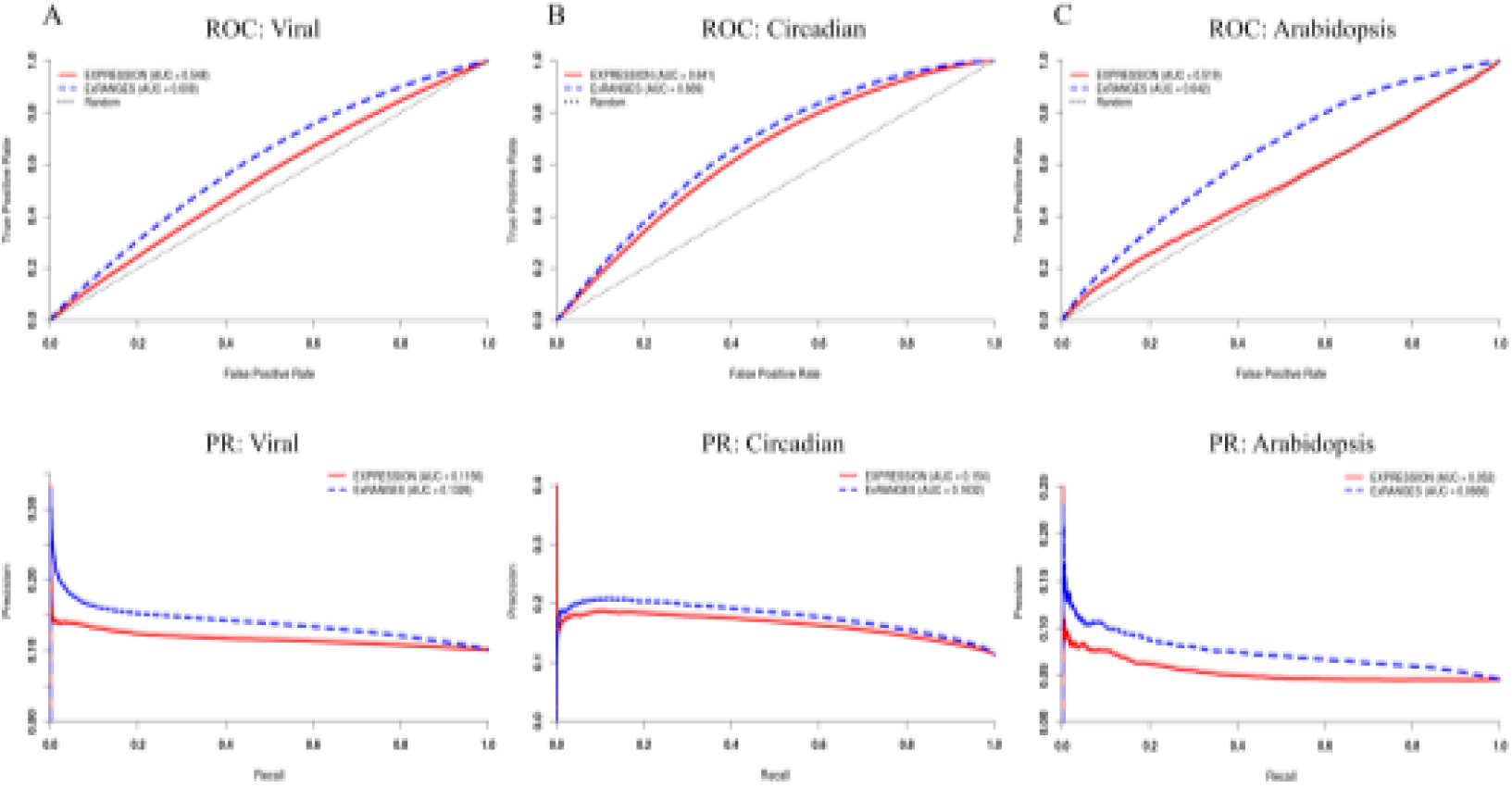
Summary of ExRANGES Improvement across Three Data Sets from Different Species. ROC and Precision-recall (PR) curves for targets of all ChIP-Seq validated TFs as identified using GENIE3 with either EXPRESSION (solid) or ExRANGES (dotted) for A) CircaDB data set from mouse tissues B) Human viral data set C) Arabidopsis circadian data set across different environmental variables.

To determine the effectiveness of the ExRANGES approach for experiments with limited time steps, we evaluated the targets identified by ExRANGES and EXPRESSION for a single time series consisting of 32 samples from eight unevenly sampled time points of field-grown rice panicles. ChIP-Seq with replicates has only been performed for one transcription factor in rice, OsMADS1 ^57^. Therefore, we compared the ability of ExRANGES and EXPRESSION to identify the OsMADS1 targets identified by L. Khanday et al. Of the 3112 OsMADS1 targets identified by ChIP-Seq, ExRANGES showed an improved ability to identify these targets (Fig. 7) compared to EXPRESSION alone.

**Figure 7:**
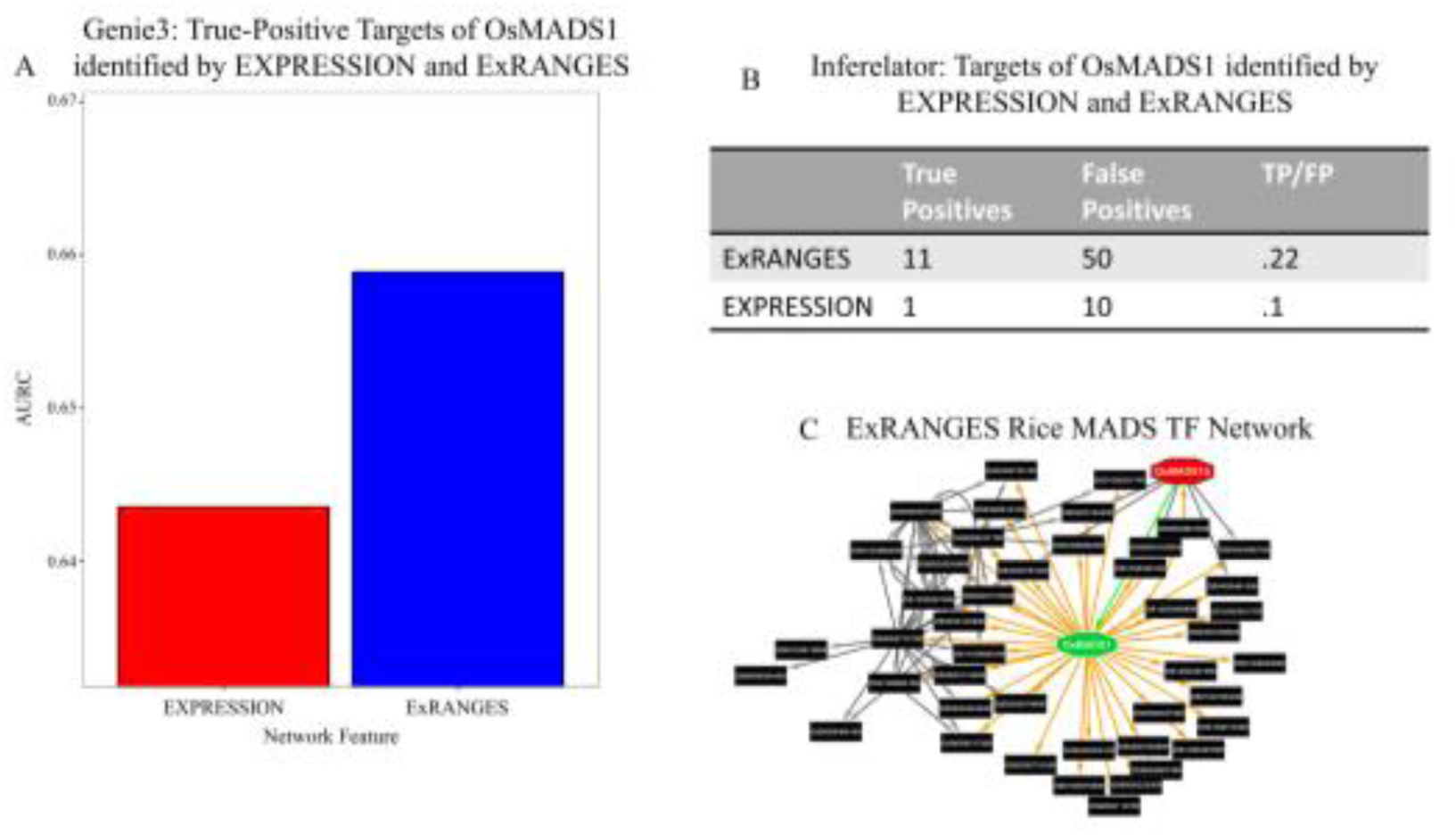
ExRANGES Retains Performance Improvement over EXPRESSION on Small Data Sets. A) ROC AUC for the top 1000 targets of OsMADS1 identified by GENIE3 using EXPRESSION or ExRANGES and validated against the OsMADS1 ChIP-Seq data ^57^. B) Comparison of targets identified by EXPRESSION and ExRANGES using INFERELATOR. ExRANGES scores higher in the ratio of True Positive (TP) to False Positives (FP). C) Interactions predicted by ExRANGES of OsMADS1 (center, green) with other MADS TFs. Orange arrows indicate ExRANGES predicted targets of OsMADS1. ExRANGES predicts that OsMADS15 (red) regulates OsMADS1 (green arrow). Interactions between other MADS TFs predicted by ExRANGES are indicated by black arrows.

## Discussion

Computational approaches that identify TRN can advance research. Most current approaches to elucidate TRN from transcriptional data use expression levels alone. We demonstrate that combining the expression levels and rate of change improves the ability to predict true targets of TFs across a range of species and experimental designs. The improvement is observed across many different collections of time series data including experiments with replicates and without, evenly and unevenly sampled time points, and even for time series with a limited number of samples. ExRANGES improves TF target identification over EXPRESSION values alone for time series performed with both microarray and RNA-Seq measurements of expression.

In many species the majority of transcripts show variation in expression levels throughout the day ^27, 47, 58^, therefore circadian and diel data sets provide a snapshot of the potential ranges in expression that a regulator can attain. Mining this daily variation can identify regulatory relationships, including those that are enhanced in response to environmental perturbations such as stress. However, one challenge with analysis of daily expression changes is that when combining multiple data sets, the daily variation in expression may be dwarfed by the large variation in expression between tissues. TRN inference approaches, such as ExRANGES that can detect the small changes in daily expression amidst the large variations between tissues are needed to fully mine this data. Here, we show that using ExRANGES, data sets that combine circadian time series in multiple tissues can be a powerful resource for identifying regulatory relationships between TFs and their targets not just for circadian regulators, but also for regulators that are not components of the circadian clock. Using EXPRESSION as the feature focused on identifying TF targets with a large variance between tissues, while targets identified using rate change showed larger variance within each time series (Fig. 2, and Supplemental Fig. 5). ExRANGES takes advantage of both sources of variation and improves the identification of TF targets for most regulators tested, including for TF-target relationships in tissues not included in the transcriptional analysis.

As implemented, ExRANGES improves the ability to identify regulator targets, however, there are many aspects that could be further optimized. For example, we tested ExRANGES with the network inference algorithms GENIE3 observed improved performance with this algorithm. ExRANGES can be applied to most other network inference algorithms. For example, we also compared the performance of Inferelator ^4^. We observe an improvement when using ExRANGES as an input with Inferelator over using EXPRESSION values alone for the viral, arabidopsis, and rice data sets (Supplemental Fig. 9 and Fig. 7B). We anticipate that ExRANGES can be integrated into other machine learning applications such as Bayesian networks, mutual information networks, or even supervised machine learning tools. Conceptually, our method increases the value of the time point before a major change in expression level. ExRANGES could be further modified to adjust where that weight is placed, a step or more in advance, depending on the time series data. Such incorporation of a time delay optimization into the ExRANGES approach could lead to further improvement for identification of some TF targets, although it would increase the computational cost.

We compared ExRANGES based features to EXPRESSION based features by validating against TF targets identified by ChIP-Seq and ChIP-Chip. While these experimental approaches identify potential TF targets in a genome-wide manner, systemic bias in ChIP could bias the comparisons^59^. For example, we observed that ChIP-seq identified targets in the CircaDB data set showed lower variation in expression than computationally identified targets (Fig. 2). The use of ExRANGES as a network input also outperformed the use of EXPRESSION alone when validated against DAP-Seq, and protein binding microarray. Even though ChIP-Seq is the gold-standard for benchmarking computational approaches to identifying TF targets, high-quality ChIP-Seq data is not available in most organisms for more than a handful of TFs. This lack of experimentally identified targets is a severe hindrance to advancing research in these species. New experimental approaches such as DAP-Seq may provide alternatives for TF target identification in species recalcitrant to ChIP-Seq analysis ^53^. Additionally, O’Malley et al. improved their recall of ChIP-Seq identified targets by selecting targets that were also supported by DNase-Seq sensitivity assays ^60, 61^. Likewise, distinguishing between direct and indirect targets predicted computationally could be enhanced by incorporation of DNase-Seq or motif occurrence information for the targets. Incorporation of such a priori information on regions of open chromatin and occurrence of cis-regulatory elements leads to improved network reconstruction ^11, 62^. Combining these integrated approaches with ExRANGES could lead to further improvements in TRN identification. Although approaches such as DAP-Seq are more global in analyses than individual ChIP-Seq assays, these genome-wide approaches require a significant investment from the community in the development of an expressed TF library collection. Integrating community-acquired experimental data with network inference approaches has been successfully applied to the *Corynebacterium* genus and pathogenic *Escherichia coli*^12–14^. In these microorganisms, a database of community provided regulatory content has enabled genome-wide predictions of regulatory interactions in novel conditions. Tools to apply these resources closely related non-model species have been effective at extending the impact of the research in these model organisms ^15–19^. For non-model systems, without such resources, computational identification of TF targets can provide an economical first pass that can be followed up by experimental analysis of predicted targets, accepting the fact that there will be false positives in the validation pipeline. In this strategy, a small improvement in the ability to identify true targets of a given TF can translate into a reduced number of candidates to test and fewer experiments that must be performed. While experimental detection of the direct targets of a given TF provides the best evidence for a TRN, we hope that the improvements provided by the ExRANGES approach can facilitate research in species where experimental identification of TF targets is experimentally challenging. ExRANGES demonstrates that consideration of *how* expression data is incorporated can contribute to the success of TRN reconstruction. We hope that this analysis will stimulate evaluation of new approaches that use alternative methods to incorporate time signals into regulatory network analysis. We anticipate that further optimization and methods for integrating expression information will lead to improvements in TRN reconstruction that will ultimately accelerate biological discovery.

## Methods

### Identifying consecutive time points with significant changes in expression

Overview: We first determine the significance of the change in expression between two consecutive time points on a per gene basis. For each genei, the background variance is derived from the change in expression of genei at all consecutive time steps in all samples across from a given data set. The change in expression between two consecutive time points is evaluated against this background and the significance is calculated (Supplemental Fig. 1). For example, the mammalian circadian data set available from CircaDB ^27^ consists of time series experiments from 12 different tissues, sampled every 2 h for 48 h (288 samples). The change in expression levels for genei between time *t* and time *t+1* was determined for each consecutive time point. Since this data is cyclical, the interval between the last time point and the first time point is also included. For the CircaDB data set, the background of each consecutive time interval across the entire time series consists of 288 slopes (12 tissues x 2 h for 48). At each time step, t the slope between *t* and *t+1* was compared to a bootstrapped version of this background generated by sampling 10,000 times with replacement. For each gene, the resulting p-value was calculated by using an empirical cumulative distribution function from the R stats package. This p-value was transformed to the –log_w_ and the sign of the change in slope was preserved (R script provided). This significance of the change at each time interval is the rate change or “RANGES” value.

### Combining EXPRESSION and Rate Change using ExRANGES

ExRANGES adjusts the expression level at each time point by multiplying the Expression level at time *t* with the significance of the change in expression, or RANGES value, from time *t* to *t+1* (Supplemental Fig. 1B). This ExRANGES value was used in lieu of the expression level to generate a TRN using GENIE3 or INFERELATOR as described below ^4, 8^.

LS is a (gene x time) matrix representing a time series experiment with genes *g* and time points *T*_(*t*=1…*N*)_:

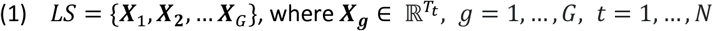

***X**_g_* is a vector of real numbers representing the expression of gene *g* from time points *T*_1_ to *T_N_*:

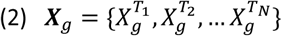

Therefore 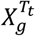 represents the expression of gene *g* at time point *T_t_*. To calculate the rate of change for RANGES values, we start with *C*_g_, which represents the changes between all consecutive time points for gene *g:*

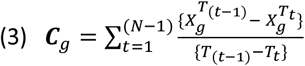

If the data are cyclical, we assume time point 1 can be used as time point N+1:

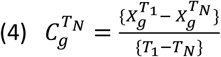

Else, disregard 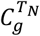

The sign of each change is recorded for use in the final RANGES value:

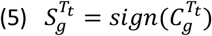

A bootstrapped version of *C_g_* is calculated for each gene by sampling 10,000 times with replacement. We call this 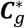. A cumulative distribution function is found for each 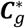.

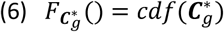

*P*-values are determined for each 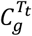 using the corresponding cumulative distribution function for each gene:

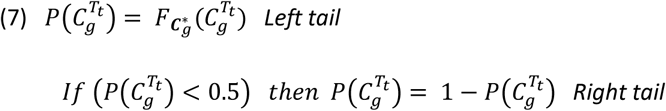

The RANGES value is calculated by taking the -log_10_ of the p-value and multiplying by the sign of the corresponding change:

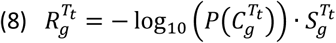

Finally, the ExRANGES value for each data point is calculated by multiplying the RANGES value by the original expression value:

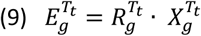

### Network Inference using GENIE3

To predict regulatory interaction between the transcription factor and the target gene, GENIE3 (http://www.montefiore.ulg.ac.be/∽huynh-thu/software.html on June 14,_2016 ^8^) was modified for use with *parLapply* from the R parallel package^63^. The EXPRESSION network was built by providing the expression values across all samples for both TFs and targets. The ExRANGES network used the ExRANGES value for both TFs and targets. For example, for the CircaDB data, we considered 1690 murine TFs as the regulators ^64^. For both approaches, all TFs were also included in the target list to identify regulatory connections between TFs. To implement GENIE3, we used 2000 trees for random forest for all data sets except the viral data set. Due to the size of the data set, we limited the viral data set to 100 trees. The importance measure from the random forest was calculated using the mean decrease in accuracy upon random permutation of individual features. This measure is used as the prediction score for TF-target relationships.

### Network Inference using INFERELATOR

For INFERELATOR the TF and targets labels are identical to those used in GENIE3. Time information in the form of the time step between each sample was added to satisfy time course conditions as a parameter, default values were used for all other parameters. Only confidence scores of TF-target interactions greater than 0 were evaluated against ChIP-Seq standards. The confidence scores were used as the prediction score for TF-target relationships.

### ROC Calculation

ROC values were determined by the ROCR package in R^65^. The computationally determined prediction score and the targets from the respective experimental validation (ChIP-Seq, protein binding array, or DAP-Seq) were used as the metric to evaluate the performance function. The area under the ROC curve (AUC) is presented to summarize the accuracy.

## Acknowledgements

We would like to thank Dahlia Nielsen, Katie Greenham, and Erin Slabaugh for critical suggestions on the manuscript preparation. Additionally, we thank Steve Briggs for sharing the time, expertise, and helpful discussions of his research group. This is contribution no. 17–389-J of the Kansas Agricultural Experiment Station. This project was supported by the Agriculture and Food Research Initiative grant # 2015–67013-22814 of the USDA National Institute of Food and Agriculture and the USDA National Institute of Food and Agriculture project 1002035.

## Author contributions statement

All authors conceived the design of the approach and J. S. D., R. C. S., and C. J. D. analyzed results and evaluated validation approaches. J. S. D., L. M. L., and S. K. J. generated *Oryza* diel time course data. C. J. D. wrote manuscript. All authors edited and reviewed the manuscript.

## Additional information

All data and scripts are either taken from existing public data or are available (Accession number: GSE92302, sources of existing data are provided in supplemental materials and methods); Competing financial interests: None declared. The corresponding author is responsible for submitting a competing financial interests statement on behalf of all authors of the paper. This statement must be included in the submitted article file.

